# Hearing scenes: A neuromagnetic signature of perceived auditory spatial extent

**DOI:** 10.1101/061762

**Authors:** Santani Teng, Verena Sommer, Dimitrios Pantazis, Aude Oliva

## Abstract

Perceiving the geometry of surrounding space is a multisensory process, crucial to contextualizing object perception and guiding navigation behavior. Auditory cues are informative about the shape and extent of large-scale environments: humans can make judgments about surrounding spaces from reverberation cues. However, how the scale of auditory space is represented neurally is unknown. Here, by orthogonally varying the spatial extent and sound source content of auditory scenes during magnetoencephalography (MEG) recording, we report a neural signature of auditory space size perception, starting ~145 ms after stimulus onset. Importantly, this neuromagnetic response is readily dissociable in form and time into representations of the source and its reverberant enclosing space: while the source exhibits an early and transient response, the neural signature of space is sustained and independent of the original source that produced it. Further, the space size response is robust to variations in sound source, and vice versa. The MEG decoding signal was distributed primarily across bilateral temporal sensor locations, significantly correlated with behavioral responses in a separate experiment. Together, our results provide the first neuromagnetic evidence for a robust auditory space size representation in the human brain, sensitive to reverberant decay, and reveal the temporal dynamics of how such a code emerges over time from the transformation of complex naturalistic auditory signals.

## Introduction

Imagine walking into a cathedral at night: Even in darkness, the passage from the narrow entryway to the large nave is immediately apparent. The reverberations produced by multiple echoes of footfalls, speech, and other ambient sounds produce a percept of the extent, or size, of each space.

Perceiving the properties of surrounding space is crucial for effective interaction with a multisensory environment [1]. Visual space representation provides context for object perception [2,3] and spatially oriented behavior such as navigation [4,5], and is mediated by similar brain structures across species [6–10]. Recent work in spatial scene processing indicates that visual environments are represented along separable and complementary dimensions of spatial boundary and content [7,11–14]. In this way a scene may be characterized by, e.g., its encompassing shape and size, as well as by the number, type, and configuration of objects it contains [15,16].

Much prior work in audition has investigated the spatial localization and perceptual organization of sound sources [17–20]. The extent to which the context, i.e. spatial environments, of these sounds, has been considered, has tended to be limited to the way reverberations from interior surfaces modulate sound-source perception. The auditory system typically works to counteract the distorting effects of reverberant environments on perception, facilitating perceptual robustness of, e.g, stimulus spatial position [21–23], speaker identity [24,25], or estimated loudness [26,27]. Yet beyond being an acoustic nuisance to overcome, reverberations themselves provide informative cues about the properties of the environment [28], and humans are able to perceive those cues to estimate features such as sound source distances [29–32] and room sizes [33–38]. Still, the auditory parameters of room size are not well understood, with nonlinear and complex relationships between physical and perceptual attributes of the auditory environment [33,34,37]. Thus, the fundamental question of how environmental properties such as space size are represented by the auditory system is generally not well understood, and its neural basis almost totally unexplored.

Here, to investigate the auditory coding of environmental space size in the human brain, we recorded magnetoencephalography (MEG) responses to auditory stimuli of sounds in different-sized spaces. We operationalized space size as the auditory room impulse response (RIR) of real-world spaces of different sizes. Auditory stimuli were constructed by convolving brief anechoic impact sounds with the RIRs, allowing us to vary scene boundary and content independently. We hypothesized that both the sound source and the size of the space could be separably decoded from the neural responses to naturalistic reverberant sounds. We found that neuromagnetic responses to spatialized sounds were readily dissociable into representations of the source and its reverberant enclosing space, and that these representations were robust to environmental variations. Our MEG results constitute the first neuromagnetic marker of auditory spatial extent processing, dissociable from sound-source discrimination, suggesting that sound sources and auditory space are processed discretely in the human brain.

## Results

To examine the neural representations of space size and auditory source information, we recorded MEG data while participants (*N* = 14) listened to three brief ~176 ms impact sounds (hand pat, pole tap, ball bounce), convolved with three room impulse responses (RIRs) corresponding to real-world indoor spaces of approximately 50, 130, and 600 m^3^ volume. The resulting stimuli were sounds spatialized to be perceived as occurring in a small, medium, or large space. Thus, three sound sources and three RIRs produced a total of nine different stimulus conditions. Participants performed a vigilance task in which occasional deviant sounds (human speech syllables) prompted a button press; these trials were excluded from analysis.

We extracted peristimulus MEG time series from -200 to +1000 ms relative to stimulus onset, and applied a linear support vector machine (SVM) at each time point to decode conditions. Statistical significance of decoding accuracy time courses were computed using one-sample t-tests against 50%, and corrected for multiple comparisons across time points using a false discovery rate (FDR) of 5% [39]. We report decoding significance onset and peak latencies with ± standard error computed by bootstrapping participants.

### Auditory representations discriminated scene size and sound source with temporally dissociable decoding trajectories

We applied SVM to decode every pair of conditions (Fig 1B) [40–42]. The pairwise MEG decoding accuracies at each time point were arranged in a 9 × 9 decoding matrix, termed *representational dissimilarity matrix* (RDM), that serves as a higher-order distance measure between conditions [43]. We then computed a single-sound decoding time course by averaging all RDM elements per time point (Fig 1C, shaded partition). The single-condition classification time course increased sharply shortly after stimulus onset, reaching significance at 60 ± 23 ms and peaking at 156 ± 20 ms. These results indicate that the MEG signal was able to reliably distinguish between individual stimulus conditions.

**Figure 1.**
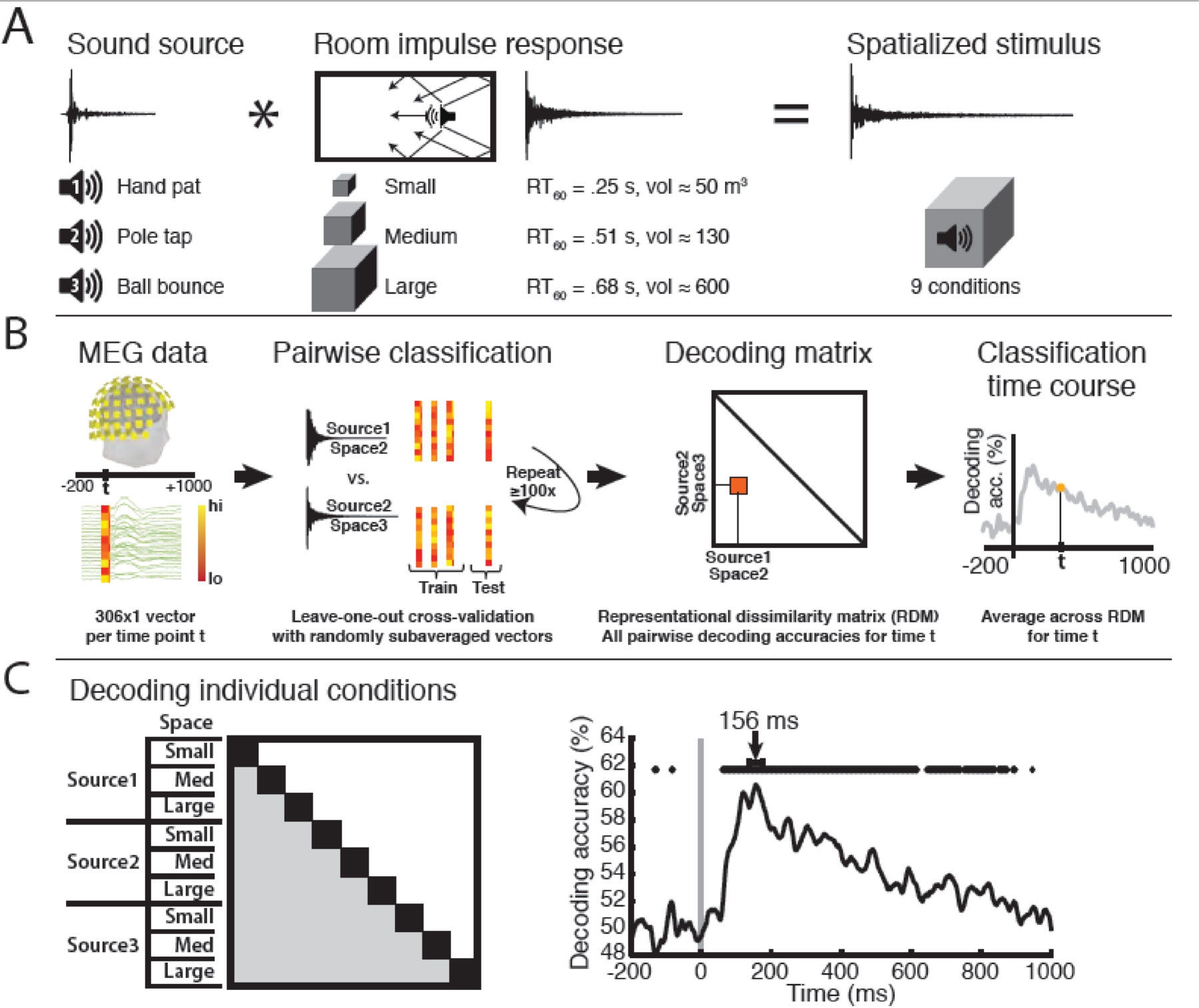
Stimulus conditions, MEG classification scheme, and single-sound decoding time course. **A**. Stimulus design. Three brief sounds were convolved with three different room impulse responses to produce nine sound sources spatialized in reverberant environments. **B**. MEG pattern vectors were used to train an SVM classifier to discriminate every pair of stimulus conditions (3 sound sources in 3 different space sizes each). Decoding accuracies across every pair of conditions were arranged in 9=9 decoding matrices, one per time point *t*. **C**. Averaging across all condition pairs (shaded matrix partition) for each time point *t* resulted in a single-sound decoding time course. Lines above time course indicates significant time points (N=14, one-sample t-test against 50%, p<0.05, false discovery rate corrected). Decoding peaked at 156 ms, with latency error bars indicating standard deviation computed by bootstrapping participants.

To dissociate the neuronal dynamics of space size and sound source discrimination, we repeated this analysis, but pooled trials across the corresponding conditions before decoding. This resulted in 3×3 RDM matrices (Fig. 2A), and averaging across the shaded regions produced the time courses of space size (red) and source identity (blue) decoding (Fig 2B). The transient nature of source discrimination, reaching significance at 59 ± 19 ms and peaking at 130 ± 10 ms, is in sharp contrast to the slower, sustained response of the space size decoding time course, which exhibited a significantly later significance onset (145 ± 43 ms) and decoding accuracy peak (386 ± 46 ms; peak latency difference, *P*<.001). This suggests that sound-source information is discriminated early by the auditory system, followed by reliable space size discrimination. In a control experiment, we found that sources and spaces were still decodable when all stimuli were controlled for duration, suggesting that the timing is not solely dependent on stimulus duration (see Fig. S1, SUPPLEMENTAL INFORMATION).

**Figure 2.**
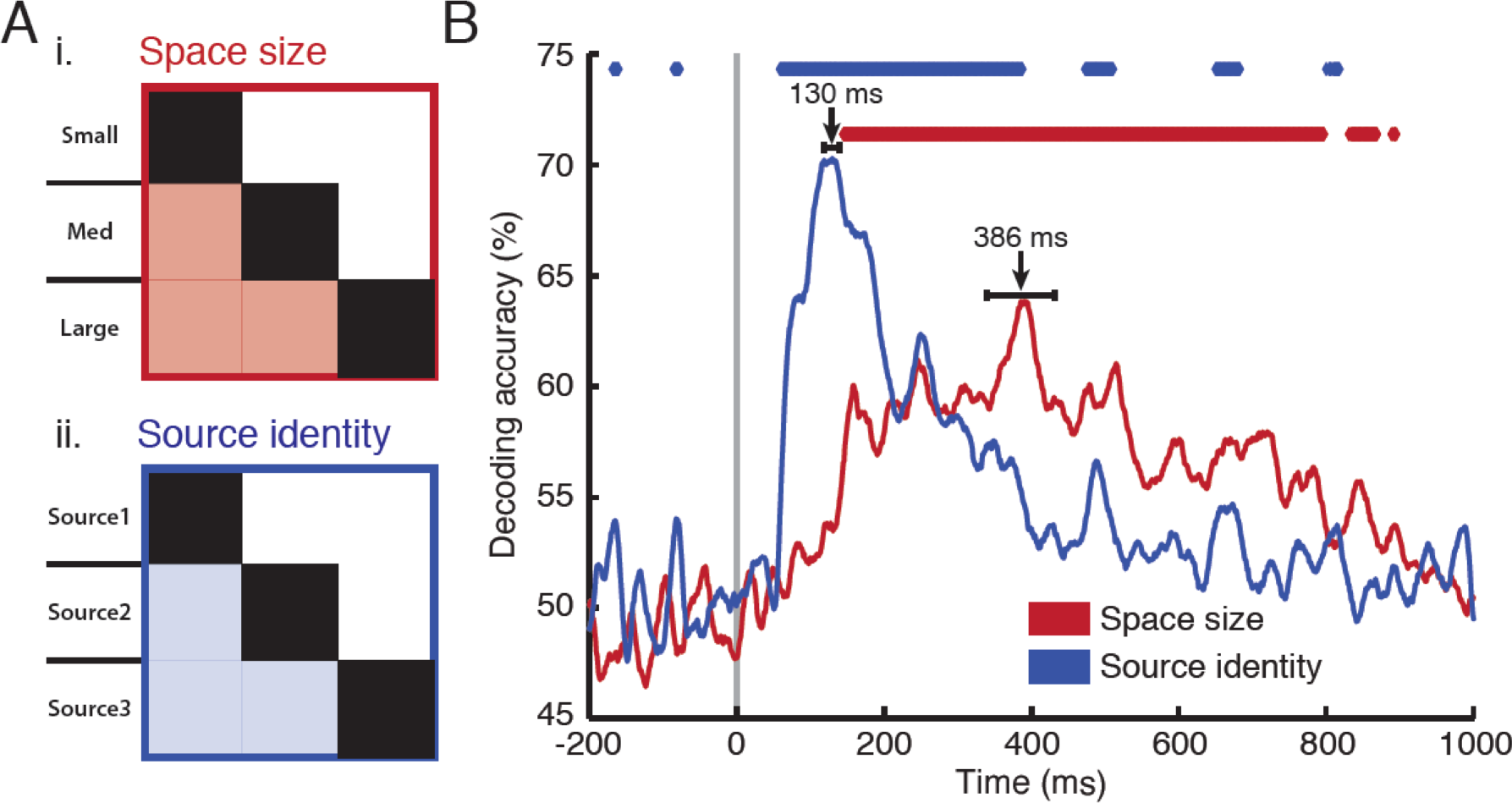
Auditory space size and source identity decoding. Individual conditions were pooled across source identity (**A***i*) or space size (**A***ii*) in separate analyses. We then performed classification analysis on the orthogonal stimulus dimension to establish the time course (**B**) with which the brain discriminated between space size (red) and source identity (blue). Sound-source classification peaked at 130±10 ms, while space size classification peaked at 386 ±46 ms. Lines above time courses and latency error bars same as in Fig. 1. See also Fig. S1 and main text.

### Space size representations are robust to variation in sound source (scene content)

Stable source and space representations should be independent of other changing properties in a scene. For example, the spatial information in a given source-space combination should be recoverable from a different source reverberating in the same space. To investigate the robustness of the respective source and space decoding curves to environmental variation, we conducted a cross-classification analysis in which we assigned space size conditions from two sources to a training set, and the size conditions from the remaining source to a testing set. Results from all such train/test combinations were averaged to produce a measure of space size information pooled across sound sources, with sound sources not overlapping between training and testing sets (Fig 3A). We then performed an analogous analysis to cross-classify sound sources. The results (Fig. 3B) indicate time courses similar to those in the pooled analysis, demonstrating that the neural representations of space size and sound source are robust to variations in an orthogonal dimension.

**Figure 3.**
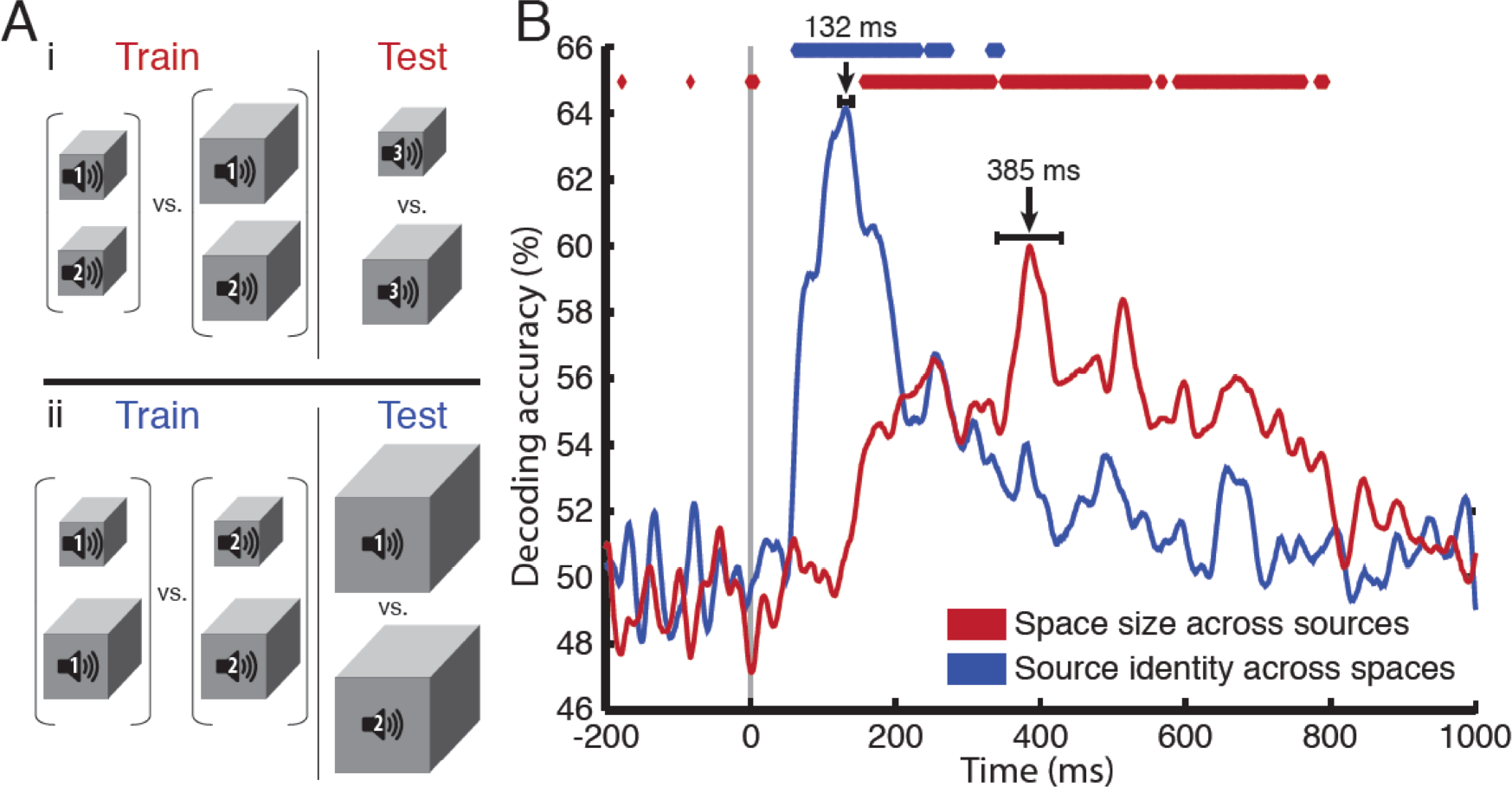
Cross-classification of space size and source identity. Space size was classified across sound sources, and sound source identity across space sizes. **A*i***. Space-size classification example in which a classifier was trained to discriminate between space sizes on sound sources 1 and 2, then tested on space discrimination on source 3. **A*ii***. Example in which a classifier was trained to discriminate between sound sources on space sizes 1 and 2, then tested on sound-source discrimination on space 3. **B**. Results from all nine such pairwise train-test combinations were averaged to produce a classification time course (**B**) in which the train and test conditions contained different experimental factors. Lines above time courses and latency error bars same as in Fig. 1.

### MEG decoding dynamics predict relative timing and accuracy of behavioral judgments

To extract behavioral parameters that could be compared with the dynamics of the MEG signal, we binned all trials into appropriate source or space comparison categories (e.g., Space1 vs. Space2; Source1 vs. Source3; etc.). Within each category we computed each subject’s mean accuracy and mean response time (mean RT estimated by fitting a gamma distribution to the response time data [44]). This yielded mean accuracies and RTs in three source-comparison and three space-comparison conditions, analogous to the pooled MEG decoding analysis. Behavioral accuracies and RTs were then correlated with MEG peak decoding accuracies and peak latencies, respectively. Significance and confidence intervals were determined by bootstrapping the behavioral and MEG subject pools 10,000 times. Behavioral RTs and peak latencies were significantly correlated (r = 0.52, p < .0072), as were behavioral accuracies and peak decoding accuracies (r = 0.57, p < .0001) (Fig. 4).

**Figure 4.**
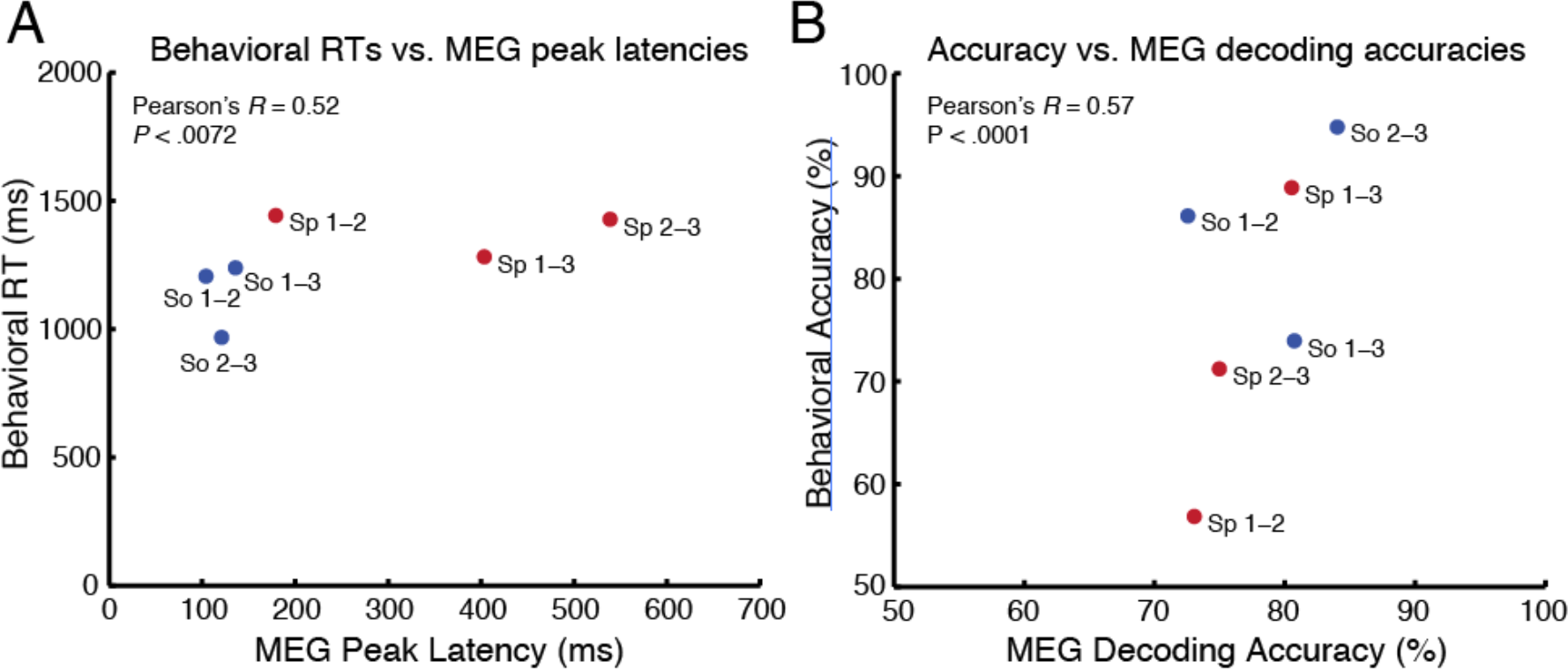
Behavioral comparison to peak decoding latency and decoding accuracy. We assessed linear relationships between response times and MEG peak decoding latencies (**A**), as well as behavioral and decoding accuracies (**B**). Bootstrapping the participant sample (N=14, p < .05) 10,000 times revealed significant correlations between RT and latency (r = 0.52, p<.0072) and behavioral and decoding accuracy (r = 0.57, p<.0001).

### Space size is encoded in a stepwise progression

The MEG space size decoding results (Fig. 2B) could be indicative of an encoded size scale (i.e., a small-to-large progression), or they could simply reflect a generic category difference between the three space size conditions. To evaluate whether MEG responses were consistent with ordinal vs. categorical size coding, we devised simple models of space representation in the form of RDMs that reflected the hypothesized space size representations. That is, each pairwise representational distance in the model 9 × 9 condition matrix was either 0 or 1, reflecting a within vs. between separation (categorical space size model), or 0, 1, or 2 reflecting a pure ordinal separation between space size conditions irrespective of sound source identity (ordinal space size model). We then correlated (using Spearman rank to capture ordinal relationships) the model RDMs with the brain response RDMs at every time point between the first and last time point of significant space decoding in the pooled analysis (145-895 ms post-stimulus onset). Fig. 5 shows that an ordinal size model correlates significantly more strongly with the ne raldata than a categorical size model (*t*-test, *P* ≪ 0.001), suggesting that space sizes have ordinal representations.

**Figure 5.**
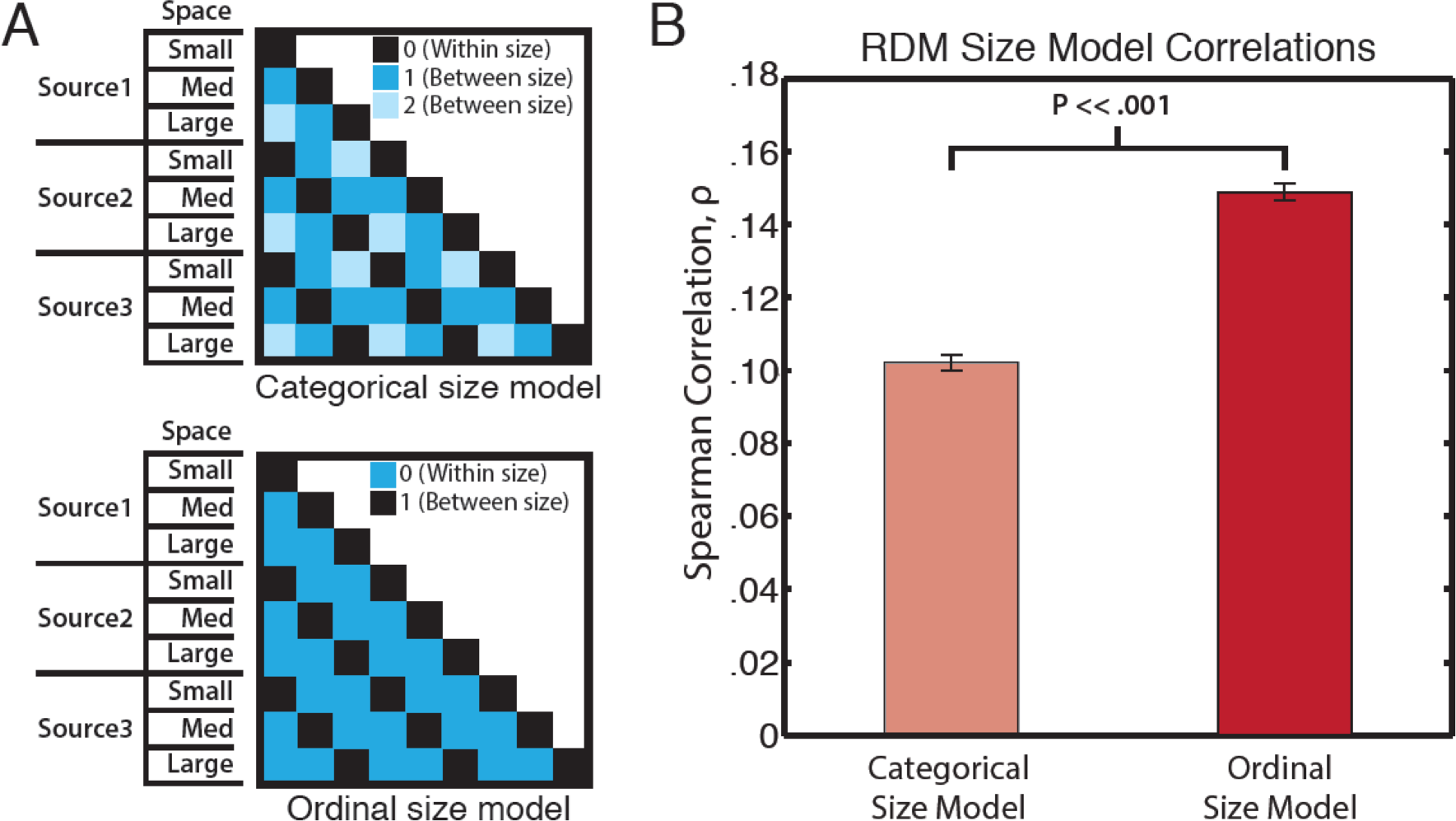
Comparison of MEG neural representations to a categorical versus an ordinal scene size model. Representational dissimilarity matrices (RDMs) of a categorical and an ordinal model (**A**) were correlated with the MEG data from 145-895 ms (the temporal window of significant space size decoding) to assess the nature of MEG scene size representations. **B**: Results indicate a significantly higher correlation between MEG representations and the ordinal size model. Spearman correlation coefficients ρ were averaged across time points in the temporal window. Error bars = ±SEM.

### Source identity and space size were best decoded by bilateral temporal sensors

To determine the spatial distribution of the decoding response, we repeated the main analysis on sensor clusters in 102 separate locations across the scanner helmet (and thus across the subject’s head). This analysis revealed that the bulk of significant decoding performance (p < .01, FDR corrected across sensors at each time point) was concentrated in sensor clusters over bilateral temporal regions (Fig. 6). While the spatial interpretation of such an analysis is limited in resolution, this result suggests that bilateral auditory cortical regions are performing the computations that distinguish among the sound sources and space sizes in the stimuli.

**Figure 6.**
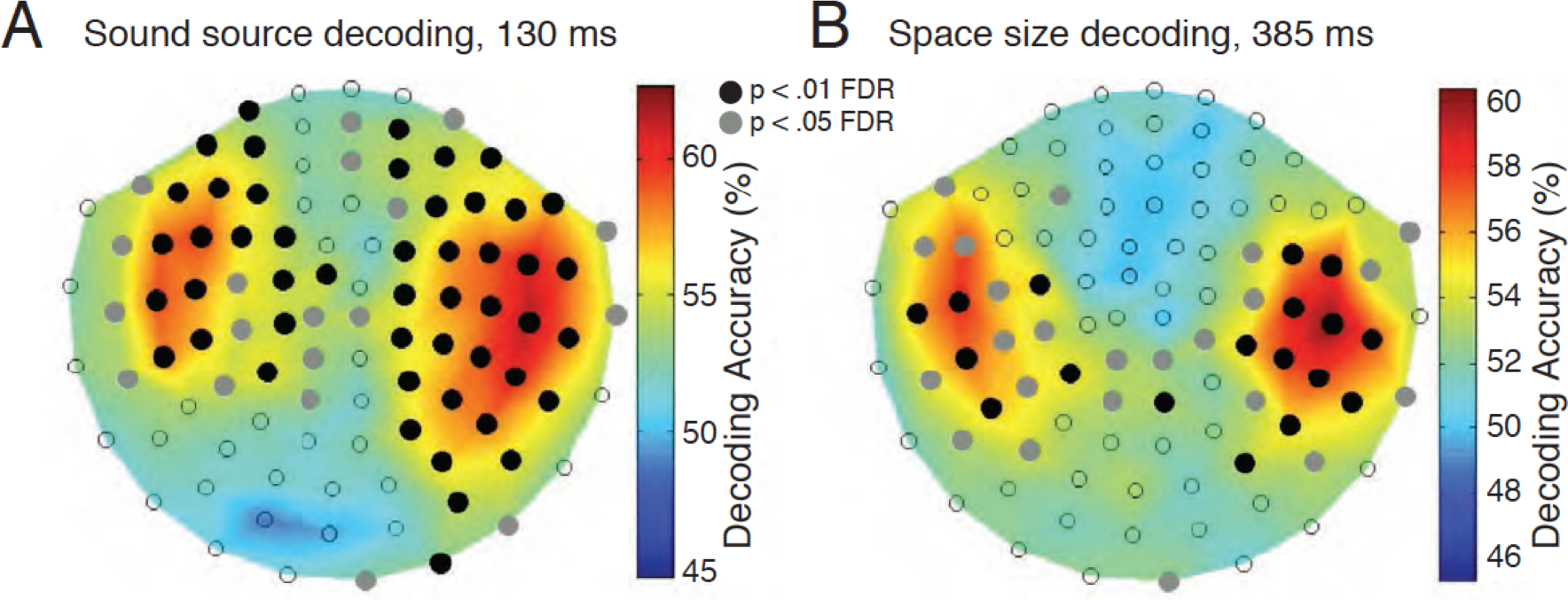
Sensorwise decoding of source identity and space size. Classification time courses were computed for each of 102 sensor triplets at each time point, corrected across sensors (and time) for a 1% FDR. Significant decoding at a given time point is indicated with a black circle over that sensor position. Times displayed are for pooled analysis decoding peaks for source identity (**A**) and space size (**B**). Orientation of sensor map is top-down, with the face pointing up.

### Dynamics of space size representations are slower and more sustained compared to sound-source representations

To examine the temporal dynamics of source and space representations, we conducted a temporal generalization analysis [41,45] in which a classifier trained at one time point was tested on all other time points. This produced a two-dimensional matrix showing generalized decoding profiles for space size and source identity (Fig. 7). The results suggest differences in processing dynamics: the narrow “diagonal chain” shown for source identity decoding in Fig. 7B indicates that classifiers trained at a time point *t* only generalize well to neighboring time points; by contrast, the space size profile (Fig. 7A) exhibits a broader off-diagonal decoding regime, indicating that classifiers were able to discriminate between space conditions over many time points other than the one in which they were trained. This suggests that space size representations are mediated by more stable, sustained underlying neural activity, compared to transient, dynamic activity mediating sound-source representations [45].

**Figure 7.**
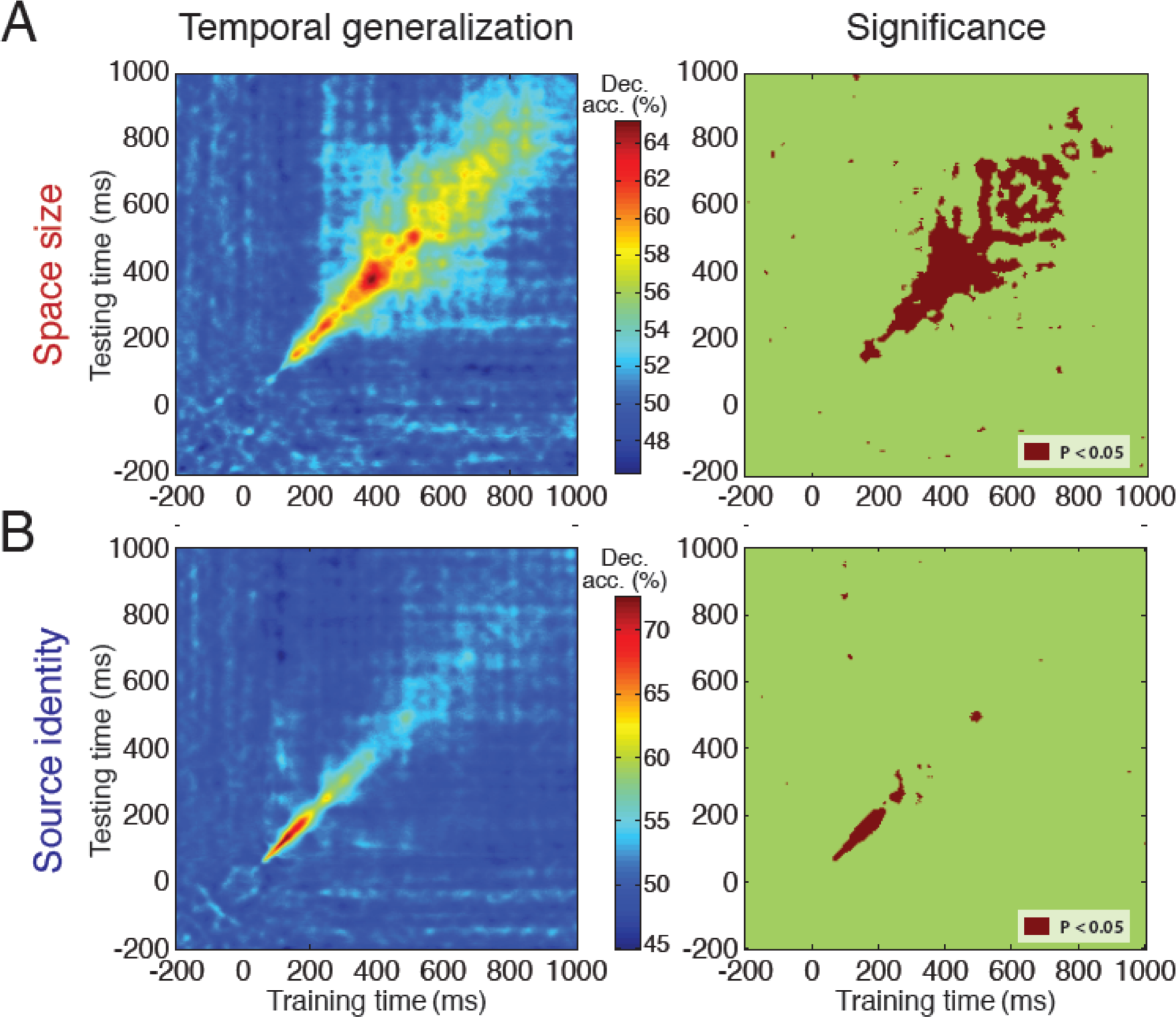
Temporal generalization matrix of auditory source and space decoding time courses. Left column plots indicate generalized decoding profiles of space (**A**) and source (**B**) decoding. Right column plots indicate statistically significant train-test decoding coordinates (*t*-test against 50%, p<0.05, FDR corrected).

## Discussion

We investigated the neural representation of auditory spatial extent and sound source identity using multivariate pattern analysis [41,42] on MEG brain responses to spatialized sounds. Our results showed that individual sound conditions were decoded starting at ~60 ms post-stimulus onset, peaking at 156 ± 20 ms. Next, we characterized the separate neural time courses of space- and source-specific discrimination. Source and space decoding profiles emerged with markedly different time courses, with the source discrimination time course exhibiting a rapid-onset transient response peaking at 130 ± 10 ms, and the space size discrimination time course ramping up more gradually to peak at 386 ± 46 ms. Further, a cross-classification analysis, in which training and testing trials contained different experimental factors, showed that the space size representation remained robust across different sounds in each space, and that the sound-source representation remained robust across different space sizes. This suggests that these representations are independent of low-level variations, such as differences in amplitude envelope and spectral distribution, that accompany environmental changes commonly encountered in real-world situations. A sensorwise decoding analysis showed that bilateral temporal cortical areas contributed most heavily to the decoding performance. Finally, the MEG decoding signal did not share the temporal profile of interstimulus correlations (Fig. S2) but was significantly correlated with behavioral responses, suggesting that the neural signal time course is (a) separate from the stimulus time course and (b) likely to reflect processing that drives perception. Taken together, the MEG decoding time courses suggest that the human auditory system extracts space size from a naturalistic stimulus.

*Dissociable and independent decoding time courses*. The pooled and cross-classified source and space decoding time courses (Figs. 2, 3) indicate that space size and sound-source representation are dissociable and independent in the brain. The decoding offset for sound source decoding suggests that source or object identity information is primarily carried by the direct sound and accompanying early reflections (the first, most nearby surface echoes [34]). By contrast, the later space size decoding peak and concurrent falloff in source decoding suggest that reverberant decay primarily carries spatial extent information. This is consistent with physical properties of large spaces, i.e. the longer propagation time for reflections to reach the listener, as well as longer RT_60_ reverberant decay times, in large spaces.

### Bilateral temporal decoding

We observed a bilateral temporal decoding response to the stimulus conditions for space sizes as well as sound source identities. While the spatial resolution of the sensorwise analysis cannot determine the exact loci of the signal sources, our results are consistent with a bilateral [46,47] account of space size processing, involving primary and nonprimary auditory cortical regions such as the planum temporale [48]. Specifically, the spatial distribution of the sensor decoding analysis argues against the interpretation that subjects were processing a related reverberation-mediated parameter such as auditory egocentric distance [49]; a previous MEG study of auditory distance reported right-lateralized sensitivity to distance in the supratemporal plane [50]. One possibility is that the spatial auditory scenes were processed by brain regions known to be selectively sensitive to visual scene size, such as the retrosplenial complex (RSC) and parahippocampal place area (PPA) [11]. However, our focus in the present study was on the dynamics of the neuromagnetic response. Localizing the exact source(s) of the space size signal is thus a target for future research.

*Relation to previous electrophysiological auditory studies*. As virtually all prior neuroimaging work on spatial auditory perception, including studies that used reverberant stimuli, measured brain responses to sound source properties, rather than properties of an enclosing space, direct comparison with previous neurophysiological results in audition is difficult. The latency of the individual- and source-decoding peaks are similar to that of numerous evoked neuromagnetic responses such as the mismatch negativity [51,50] and N1m response [52]. This generally suggests that low-level, pre-attentive evoked responses such as the P50 [50] are not strongly informative for distinguishing among the spatial conditions, but that the neural activity underlying these evoked components may be driving the general or source-specific MEG decoding performance. Later responses to naturalistic spatial stimuli include elevation-related processing starting at 200 ms [53] and a component indexing “spatiality” of sound sources, i.e. the difference between simple binaural cue manipulation and realistic 3D spatialization [54]. By contrast, the later rise of space size decoding performance in our results suggests that a different underlying mechanism is operating to code space size.

The most closely related neuroimaging investigations of auditory spatial parameters are likely those that probe the representation of egocentric distance of a sound source [31,50]. The distance to a sound source implies the minimum extent of the enclosing space, and its perception is mediated by a reverberant cue (the direct-to-reverberant energy ratio, D/R, rather than RT_60_ [29,31]). An MEG study of sound source distance found significant mismatch responses to auditory and duration deviants for reverberant virtual stimuli in data analyzed between 140 and 220 ms [50], which overlaps with the onset of the space size decoding response in the present study. If the same mechanism underlies both sets of results, the common parameter would be closer to a space size representation, as the sound sources in our stimulus set were all at the same virtual egocentric distance from the listener.

Taken together, comparison with previous work suggests that while the same substrates processing 2-D sound source location (azimuth and elevation) are unlikely to also process spatial extent of auditory scenes, the temporal dynamics of brain responses to distance may involve space size rather than object distance *per se*.

*The metric of space size representation*. Finally, while behavioral room size judgments have been previously shown to be driven by reverberant information, the precise relationship between volumetric spatial extent, reverberation time, and perceived size is nonlinear and not fully understood [33,34,29]. We used a well discriminable sequence of space sizes to establish an ordinal representation, but future work with condition-rich designs can more precisely characterize the metric of neural representation of auditory spatial extent. Further, given the parallels with the dissociable encoding of properties of visual scenes [12], future work may elucidate whether spatial extent coding shares a neural mechanism across visual and auditory sensory modalities.

In sum, the current study presents the first neuromagnetic evidence for an auditory scene size representation in the brain. The neurodynamic profile of the processing stream is dissociable from that of sound sources in the scene, robust to variations in those sound sources, and predicts both timing and accuracy of corresponding behavioral judgments. Our results establish an auditory basis for investigations of scene processing, suggest the spatial importance of the reverberant decay in perceived scene properties, and lay the groundwork for future auditory and multisensory studies of perceptual and neural correlates of environmental geometry.

## Materials and Methods

### Participants

We recruited 14 healthy volunteers (9 females, age mean ± s.d. = 27.9 ±5.2 y) with selfreported normal hearing and no history of neurological or psychiatric disease. Participants were compensated for their time and provided informed consent in accordance with guidelines of the MIT Committee on the Use of Humans as Experimental Subjects (COUHES).

### Stimuli

Stimuli were recordings of three different brief monaural anechoic impact sounds (hand pat, pole tap, and ball bounce), averaging 176 ms in duration. Each sound was convolved with three different monaural room impulse responses (RIRs) corresponding to real-world spaces of three different sizes, yielding a total of nine spatialized sound conditions. The RIRs were measured by recording repeated Golay sequences broadcast from a portable speaker and computing the impulse response from the averaged result. The speaker-recorder separation was constant at ~1.5 m. Room sizes corresponded to everyday spaces, with estimated volumes (based on room boundary dimensions) of approximately 50, 130, and 600 m^3^. Reverberation times (RT_60_, the time for an acoustic signal to drop by 60 dB) of the small, medium, and large-space RIRs were 0.25 s, 0.51 s, and 0.68 s, respectively. RT_60_ estimates given in Fig. 1 were averaged across all frequencies from 20 Hz to 16 kHz, logarithmically weighted. Auditory stimuli are available as supplemental material (see Table S1).

### MEG testing protocol

We presented stimuli to subjects diotically through tubal-insert earphones (Etymotic Research, Elk Grove Village, IL, US) at a comfortable volume, approximately 70 dB SPL. Stimulus conditions were presented in random order with stimulus onset asynchronies (SOAs) jittered between 2000-2200 ms. Every 3 to 5 trials (4 on average), a deviant vigilance target (brief speech sound) was presented, prompting participants to press a button and blink. SOAs between vigilance target and the following stimulus were 2500 ms. Target trials were excluded from analysis. Each experimental session lasted approximately 65 min and was divided into 15 runs containing 10 trials from each condition, for a total of 150 trials per condition in the entire session.

### Behavioral testing protocol

In our MEG scanning protocol, we used a passive-listening paradigm to avoid contamination of the brain signal with motor-response artifacts. Thus, to test explicit perceptual judgments of the auditory scene stimuli, we conducted separate behavioral tests of space size and sound source discrimination. Participants (N=14) listened to sequential pairs of the stimuli described above, separated by 1500 ms SOA. In separate blocks, participants made speeded same-different judgments on the sound sources or space sizes in the stimulus pairs. Condition pairs and sequences were counterbalanced for each subject, and the order of source- and space-discrimination blocks was counterbalanced across subjects. Over the course of four blocks lasting a total of ~40 min, participants completed a total of 36 trials per category. We collected reaction time and accuracy data from participants’ responses.

### MEG data acquisition

MEG recordings were obtained with an Elekta Neuromag TRIUX system (Elekta, Stockholm, Sweden), with continuous whole-brain data acquisition at 1 kHz from 306 sensors (204 planar gradiometers; 102 magnetometers), filtered between 0.3 and 330 Hz. Head motion was continuously tracked through a set of five head-position indicator coils affixed to the subject’s head.

### MEG preprocessing and analysis

Data were motion-compensated and spatiotemporally filtered offline [55,56] using Maxfilter software (Elekta, Stockholm, Sweden). All further analysis was conducted using a combination of Brainstorm software [57] and Matlab (Natick, MA, US) in-house analysis scripts. We extracted epochs for each stimulus presentation with a prestimulus baseline of 200 ms, and 1000 ms post-stimulus onset, removed the baseline mean from each sensor, and applied a 30-Hz low-pass filter.

### MEG multivariate analysis

To determine the time course of auditory space size and object identity discrimination, we analyzed MEG data using a linear support vector machine (SVM) classifier ([58]; libsvm: http://www.csie.ntu.edu.tw/~cjlin/libsvm/). For each time point *t*, the MEG sensor data were arranged in a 306-dimensional pattern vector for each of the M=150 trials per condition (Fig. 1B). To increase SNR and reduce computational load, the M single-trial pattern vectors per condition were randomly subaveraged in groups of k=10 to yield M/k subaveraged pattern vectors per condition. We then used a leave-one-out crossvalidation approach to compute the SVM classifier performance in discriminating between every pair of conditions. The whole process was repeated K=100 times, yielding an overall classifier decoding accuracy between every pair of conditions for every time point *t* (Fig. 1B).

The decoding accuracies were then arranged into 9×9 *representational dissimilarity matrices* (RDMs; [43]), one per time point *t*, indexed by condition and with the diagonal undefined. To generate the single-sound decoding time course (Fig. 1C), a mean accuracy was computed from the individual pairwise accuracies of the RDM for each time point.

For space size decoding (Fig. 2B), conditions were pooled across the three sound-sources, resulting in 3M trials for each space size. An SVM classifier was trained to discriminate between every pair of space sizes and results were averaged across all pairs. Decoding procedures were similar as above, but subaveraging was increased to k=30 and repetitions to K=300. For sound-source decoding (Fig. 2B), we pooled across the three space sizes and performed the corresponding analyses.

Statistically significant time points were determined via random-effects analysis (one-sample *t*-test against 50%, p < 0.05, false discovery rate (FDR) corrected across time points [39]). Latency error bars for significance onset and peak decoding were determined by bootstrapping participants 1000 times, computing onset and peak latencies for each bootstrap sample, and estimating the standard deviation of the resulting distribution. The standard errors of the mean latencies are depicted graphically in the figures and described in the text using the ± notation.

### MEG spatial (sensorwise) analysis

To characterize the spatial distribution of the decoding time course, we conducted a sensorwise analysis of the MEG response patterns. Specifically, the 306 MEG sensors are physically arranged in 102 clusters of three sensors each on the scanner. Thus, we performed the multivariate analysis described above at each of the 102 sensor location, using a three-(rather than 306-) dimensional pattern vector for each location. The same subaveraging and crossvalidation approach as in the main analysis was used to produce 9 × 9 RDMs of pairwise classification accuracies at each sensor position and at each time point. Thus, rather than the single whole-brain decoding time course shown in Figs. 1–3, this analysis generated 102 decoding time courses, one for each sensor cluster position. Statistically significant decoding accuracies were determined via random-effects analysis, p < 0.01, FDR-corrected across sensor positions at each time point.

### MEG temporal generalization analysis

To further interrogate the temporal dynamics of space size and source identity processing, we extended the above analysis to time-time decoding, i.e. training each classifier at a given time point *t* and testing decoding accuracy against all other time points. This yielded a 2-dimensional temporal generalization (time-time) matrix (Fig. 7) [45]. Statistical significance maps for each matrix were generated similarly to the 1dimensional analysis (*t*-test across participants for each time-time coordinate, p<0.05, FDR corrected).

### MEG cross-classification analysis

To determine the robustness of space size and sound source representations to environmental variation, we performed a cross-classification analysis in which different orthogonal experimental factors were assigned to training and testing sets. For example, the cross-classification of space size (Fig. 3A*i*) was conducted by training the SVM classifier to discriminate space size on two sound sources and testing it on the third sound source. This analysis was repeated for all such train-test combinations and the results were averaged to produce the final cross-classification accuracy plots. SVM decoding was performed similarly to the single-condition analyses, but the training set had 2M trials, subaveraging was set to k=20, and repetitions to K=150. Cross-classification of sound source identity across space sizes was performed with corresponding analyses.

## Author Contributions

S.T., D.P. and A.O. co-designed the experiments. S.T., V.S. and D.P. collected and analyzed experimental data. V.S. generated stimuli. D.P. and A.O. supervised data collection. All four authors contributed to writing the manuscript.

## Acknowledgments

We are grateful to James Traer for assistance in stimulus design and Radoslaw Cichy for helpful comments on earlier versions of this manuscript. This work was funded by National Eye Institute grant EY020484 (to A.O.), the McGovern Institute Neurotechnology Program (to A.O. and D.P.), and was conducted at the Athinoula A. Martinos Imaging Center at the McGovern Institute for Brain Research, MIT.

